# An interferon lambda 4-associated variant in the hepatitis C virus RNA polymerase affects viral replication in infected cells

**DOI:** 10.1101/2020.06.22.164319

**Authors:** Connor G G Bamford, John McLauchlan

## Abstract

Host *IFNL4* haplotype status contributes to the development of chronic hepatitis C virus infection in individuals who are acutely infected with the virus. *In silico* studies revealed that specific amino acid variants at multiple sites on the HCV polyprotein correlate with functional single nucleotide polymorphisms (SNPs) in the *IFNL4* locus. Thus, SNPs at the *IFNL4* locus may select variants that influence virus replication and thereby outcome of infection. Here, we examine the most significantly *IFNL4*-associated amino acid variants that lie in the ‘Lambda (L) 2 loop’ of the HCV NS5B RNA polymerase. L2 loop variants were introduced into both sub-genomic replicon and full-length infectious clones of HCV and viral replication examined in the presence and absence of exogenous IFNλ4. Our data demonstrate that while mutation of NS5B L2 loop affects replication, individual *IFNL4*-associated variants have modest but consistent effects on replication both in the presence and absence of IFNλ4. Given the strong genetic association between these variants and *IFNL4*, these data suggest a nuanced effect of each individual position on viral replication, the combined effect of which might mediate resistance to the effects of IFNλ4.

## Full-text

Clearance of HCV is associated with genetic and functional variation in the human IFN lambda 4 (*IFNL4*) gene (1). Recent analyses of unbiased ‘genome-to-genome’ variant association has also identified correlations between HCV genetic polymorphisms at specific sites across the virus genome and *IFNL4* variation (2–4). This suggests that virus populations in those producing functional *IFNL4* differ from those generating the non-functional or less potent forms of the protein. Thus, there may be an interaction between host and viral genetic variants that ultimately affects viral chronicity. A previous report characterised one variant in NS5A, which was linked to serum viral load in individuals expressing functional *IFNL4* indicating that this *IFNL4*-associated site may affect virus replication in the sub-genomic replicon (SGR) system (2). However, studies on robust full-length HCV cell culture (HCVcc) infectious systems have not been carried out. Furthermore, the most significant *IFNL4*-associated variant (A150V) in the NS5B protein has not been examined in detail for any contribution to the viral replication process in such systems.

The region encompassing A150V and an additional cluster of *IFNL4*-associated variants is located between amino acids (aa) aa2567 to aa2576 in the HCV polyprotein and lie towards the N-terminus of the virus-encoded NS5B RNA-dependent RNA polymerase (RdRp; positions 2567, 2568, 2570, 2576 correspond to residues 147, 148, 150 and 156 in NS5B; **Fig 1A**). This region corresponds to a relatively variable segment termed ‘motif F’ in the N-terminal finger domain of conserved viral RdRp enzymes of RNA viruses, and has been termed the lambda (L) 2 loop (**Fig 1A and B**) (5,6). Comparison of HCV RdRp sequences across other families in the *Flaviviridae*, including the Flaviviruses, Pegiviruses and Pestiviruses, revealed invariant amino acid residues flanking the L2 loop in regions termed F1 and F2. Upstream of the L2 loop is a conserved KXE motif (X=N/K/R depending on the virus family) while in the downstream region are invariant lysine, arginine and isoleucine residues in a KXXRXI motif (**Fig 1B**). Within the L2 loop there were no invariant residues. Sequence comparison of amino acid sequences for the L2 loop among and within HCV genotypes further demonstrated variability of this region (**Fig 1C**). Positions 148 and 150 showed the greatest variability although we noted also lack of sequence identity at positions 146 and 154. In addition, the proline residue in the F2 region displayed some intergenotype diversity. Based on the crystal structure of HCV NS5B the L2 loop corresponds to a flexible surface-exposed loop in the closed conformation of the protein where it extends outwards and over the nucleotide tunnel (**Fig 1D and E**) (6). Amino acids at positions 148 and 150 with the greatest variability lie at the extremity of the loop.

**Figure 1.**
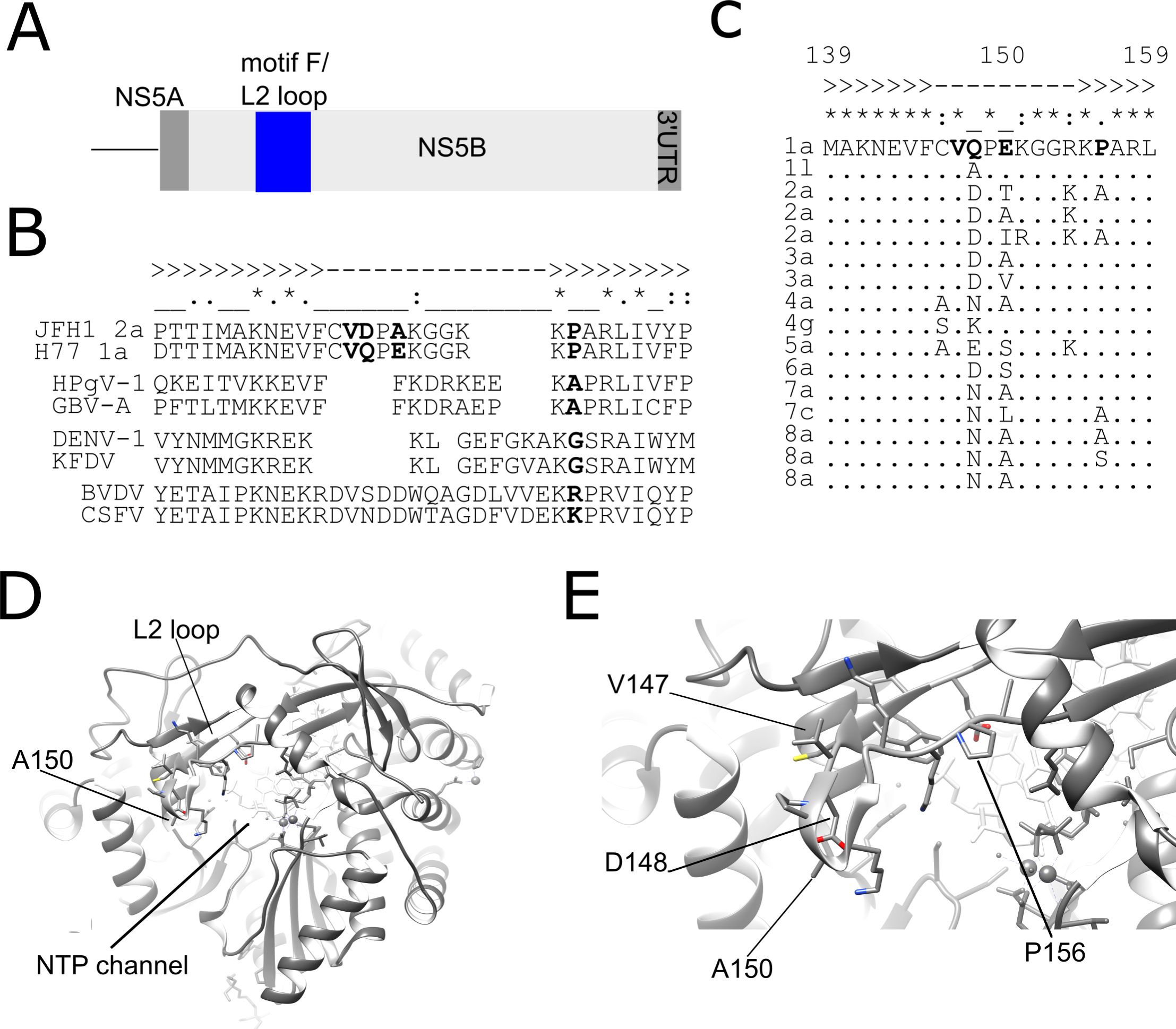
Location of *IFNL4*-associated positions in the L2 loop of HCV NS5B RNA polymerase. (A) Schematic of the 3’ end of the HCV genome showing the location of the NS5B motif F/L2 loop. (B and C) Amino acid alignment of the region encompassing motif F in the Flaviviridae (B) and in HCV reference sequences for genotypes 1-8 (C). *IFNL4*-associated positions are highlighted in bold in HCV reference sequences (for gt1a and gt2a in [B] and gt1a in [C]). Invariant (*), conserved (:) and partially conserved (.) positions are highlighted directly above the alignment. Secondary structural elements in the HCV NS5B RNA polymerase are indicated above sequences (>> for β-strands for F1 and F2; -- for the non-structured L2 loop). HCV gt2a JFH1: AB047639.1; HCV gt1a H77: AF009606.1; human pegivirus 1: BAA19580.1; Simian pegivirus/GB-virus A: AHH32939.1; Dengue virus serotype 1 (DENV-1): QFS19562.1; Kyasanur Forest disease virus (KFDV): AXB87737.1; Bovine viral diarrheal virus (BVDV): AAF82566.1; classical swine fever virus (CSFV): AYE19937.1. HCV subtype accession numbers are as follows: 1a_AF009606, 1l_KC248193, 2a_D00944, 2a_AB047639, 2a_HQ639944, 3a_D17763, 3a_JN714194, 4a_Y11604, 4g_FJ462432, 5a_AF064490, 6a_Y12083, 7a_EF108306, 7_KU861171.1, 8_pt1 MH590698.1, 8_pt2 MH590699.1, and 8_pt4 MH590701.1. (D and E) Structure of the L2 loop region in the HCV NS5B RNA polymerase. *IFNL4*-associated positions are shown with amino acid side-chains on the 4wti pdb structure of HCV gt2a JFH1 NS5B. The L2 loop is shown relative to the NTP channel and residue A150 (D) and *IFNL4*-associated positions are indicated in a higher magnification image (E).

Based on the above analyses, we constructed four SGR and HCVcc L2 loop mutants in a JFH1 gt2a SGR construct containing a GLuc reporter and the Jc1 HCVcc infectious clone (7,8). We chose to utilise the JFH1/Jc1 system because the documented *IFNL4*-associated variable position 150 is found not only in gt3a but also in gt2 sequences. Therefore we were able to exploit the high replication capability of the well-characterised JFH1/Jc1 system. Our strategy was to create substitutions at positions 148 (D148A), 150 (A150V) and a double substitution at positions 148 and 150 (D148Q.A150E) thereby reconstituting the gt1a sequences at these positions (**Fig 1C**). Gt1a is considered a relatively IFN-resistant HCV subtype and these positions have been implicated in this phenotype as well as association with IFNL4/IL28B genotype (9). We also constructed a P156A variant, identified as associated with *IFNL4* genotype in 4 HCV subtypes (2–4) to determine whether altering this position in the F2 region created a functional defect that would affect replication. Briefly the NS5B-coding region was sub-cloned into a plasmid vector and site-directed mutagenesis used to alter residues before transferring the mutated fragments into the SGR plasmid and then into HCVcc plasmids. Mutagenesis primers used are available on request. All mutations in the final constructs were confirmed by sequencing.

RNA from the SGR constructs was generated by *in vitro* transcription (IVT) using the manufacturer’s instructions (T7 RiboMAX, Promega, UK) and transfected (200ng) into sub-confluent monolayers of Huh7 cells in 96 well plates using Lipofectamine 2000 (1µl per µg RNA) using manufacturer’s instructions (ThermoFisher Scientific, UK). GLuc activity in 5% of the supernatant (10µl) was assessed at 4, 24, 48 and 72hrs post transfection using the manufacturer’s instructions. Reporter activity for each construct was compared to a replication-defective ‘GND’ mutant (**Fig 2A**). Each of the mutant constructs plateaued around 48hrs post transfection and reached a peak activity by 72hrs, achieving levels that were not significantly different from the WT construct. However, we noted that D148A gave 10-fold lower luciferase activity at 24hrs and 48hrs post transfection. Both the double mutant D148Q.A150E and A150V constructs also yielded less activity than WT (∼5-fold and ∼2-fold, respectively) at earlier times post transfection.

**Figure 2.**
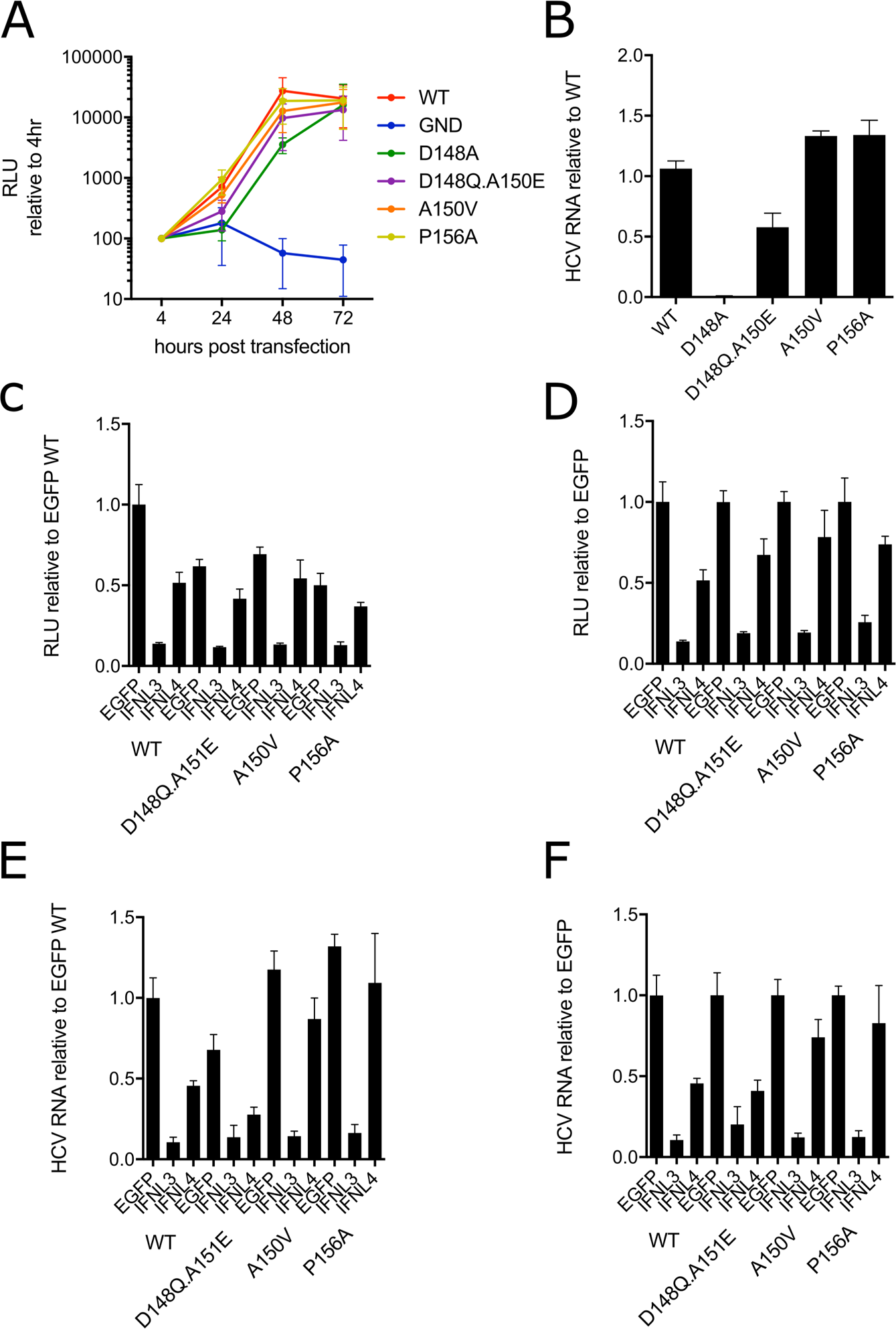
Effect of *IFNL4*-associated variants in the NS5B L2 loop on viral RNA replication in the presence and absence of IFNλ proteins. (A) Gluc activity at 4, 24, 48 and 72hrs post transfection of Huh7 cells with *in vitro* transcribed SGR RNA. SGR constructs used are indicated and include a replication-incompetent ‘GND’ control. Data are shown relative (%) to the 4 hour Gluc activity for each construct (*n*=5) (B) Viral genomic RNA abundance at 72hpi following infection of Huh7 cells at a MOI of 0.01 with the indicated WT and mutant HCVcc viruses. Data are shown relative to WT HCVcc RNA, which was normalised to 1 (*n*=2). (C and D) Sensitivity of WT and mutant SGRs (C and D) or HCVcc (E and F) to addition of exogenous IFNλ3 and IFNλ4 compared to an EGFP control. Gluc activity (C and D) or HCV genome RNA abundance (E and F) at 48hrs post transfection of Huh7 cells with *in vitro* transcribed SGR RNA or 72hpi following infection with HCVcc (MOI of 0.01) following treatment (24hrs) with IFNλ3 or IFNλ4 conditioned media (CM) (1:4) as well as the EGFP negative control CM. Data shown relative to WT SGR RNA/HCVcc from EGFP-treated cells (C and E) or to each construct EGFP-treated control (D and F) normalised to ‘1’ (*n*=2). Abundance of viral genomic RNA was measured at 72hpi relative to GAPDH mRNA. All data show mean (+/- standard error or the mean).

The same four mutants were introduced into the Jc1 HCVcc system to allow analysis of viral multicycle replication. Following IVT of HCVcc RNA and electroporation into Huh7 cells virus-containing supernatants were harvested at least 3 days later and infectivity titrated by TCID50 using an NS5A-specific antibody (10). All RNAs produced infectious virus, although D148A yielded ten-fold less infectious virus (data not shown). Sequences of the mutated versions of HCVcc were confirmed by Sanger sequencing of RT-PCR amplicons; no changes at the consensus level were found apart from D148A, which failed to yield an RT-PCR product, presumably due to reduced replication of viral RNA (data not shown). Assessment of HCVcc RNA accumulation at 72hpi following infection of Huh7 cells infected at MOI of 0.01 showed that while HCVcc viral RNA was detected by RT-qPCR as previously described (7) in all infections, only D148A and D148Q.A150E showed major differences (**Fig 2B**). D148A gave a 100-fold reduction in RNA accumulation and D148Q.A150E a more modest 2-fold reduction in RNA accumulation. A150V and P156A led to slight (less than 1.5-fold) but consistent increases in viral RNA accumulation.

One of our main objectives in the study was to examine whether A150V, which is associated with host *IFNL4* genotype, would affect multicycle HCV replication in the presence or absence of IFNλ4 using an *in vitro* model system. We have previously established a system to test the antiviral activity of WT human IFNλ3 and IFNλ4 on HCV infection (11). Huh7 cells, treated with exogenous transfected-cell ‘conditioned media’ (CM) containing IFNs, were transfected with WT and mutant SGR RNAs or infected with WT and mutant HCVcc; GLuc activity was assayed at 48hpt (for SGR) and viral RNA measured at 72hpi (for HCVcc). Given the low replication of SGR and HCVcc constructs carrying the D148A mutation, we excluded these constructs from further analysis. Pre-treatment of Huh7 cells with exogenous IFNλs gave a greater reduction with IFNλ3 compared to IFNλ4 for Gluc activity from SGR (**Fig 2C and D**) and viral RNA from HCVcc constructs (**Fig 2E and F**) compared to cells treated with a control CM from EGFP plasmid-transfected controls. These results were similar to our previous analyses (11). SGRs containing mutations had reduced replication in EGFP-CM-treated cells compared to WT (**Fig 2C**), consistent with data presented in Fig 2A at 48hpt (**Fig 2A**). IFNλ pre-treatment inhibited SGR replication and, by normalising the data to that from EGFP CM-treated cells, the mutations introduced into the SGR had a modest effect on replication in IFNλ4-treated cells, with all mutants yielding approximately ∼1.5-fold higher Gluc levels compared to the WT construct treated with IFNλ4 (**Fig 2D**). By comparison, IFNλ3 gave a more pronounced inhibition for all SGRs. Similar to the data in Fig 2B, the HCVcc D148Q.A150E mutant had reduced replication in EGFP-CM-treated cells compared to WT (**Fig 2E**). IFNλ pre-treatment inhibited HCVcc RNA synthesis and, from normalisation with EGFP CM-treated cells, showed that the D148Q.A150V mutant led to similar RNA accumulation as the WT construct (**Fig 2F**). However, the A150V and P156A substitutions yielded a ∼1.5-fold increase in RNA levels similar to what was observed in SGR assays, suggestive of a consistent lower fall in replication compared to the WT construct in the context of prior IFNλ4 treatment. Again, IFNλ3 pre-treatment robustly suppressed viral replication and we observed no apparent differences between mutants. We conclude that in the presence of exogenous IFNλ4 viral replication is modestly elevated for the A150V and P156A variants, located in the L2 loop in NS5B.

All of the variants introduced into the L2 loop and F2 region in the replicon and infectious systems were natural polymorphisms that occur in at least one HCV subtype (**Fig 1C**). Since they were natural variants, we expected that they may have modest effects on viral RNA replication and virion production. However, we observed a substantial drop in viral RNA levels at earlier times in the replicon assay and poor yields of virus for the D148A substitution. An alanine residue is found at position 148 only for gt1l, a rare subtype that is typically found in sub-Saharan Africa. By contrast, replacing both residues at positions 148 and 150 recreated the most frequently found amino acids in gt1a (D148Q.A150E) and replication was decreased by <2-fold. Moreover, the other variants that were studied (A150V and P156A) had no overall effect on replication in the SGR system when compared to the WT construct and indeed, gave a modest increase in viral RNA in the infectious model. Therefore, except for D148A, the substitutions were well tolerated in NS5B.

Recent molecular dynamics data has provided greater insight into the likely function of the HCV F domain in organising the entry of nucleotides for access to the active site as well as exit of pyrophosphate during the replication process (12). Prior to nucleotide access to the active site, the F domain coordinates nucleotide reorientation and base stabilisation through rearrangement of salt bridge interactions involving K151 (with either D352 or D387 depending on conformation) and R158 (with E143). The other residues comprising the F1-L2-F2 region are likely to provide functional properties such as coordinating the opening/closure of the nucleotide tunnel, structural flexibility and perhaps some nucleotide selectivity. Furthermore, a K151R mutation rescued infectivity of a P7 mutant HCVcc but had no measurable effect on replication nor NS5B activity *in vitro* (13). Another study identified a potential interaction between the L2 loop and domain II of NS5A (14). Thus, the L2 loop may participate in interactions with other regions of NS5B (for RNA replication) and with other viral proteins like NS5A and P7/NS2 (e.g. for assembly). Both D148 and A150 are located on the surface of L2, facing downwards towards incoming nucleotides. These residues lie immediately to the N-terminal side of residue K151, which is nearly completely invariant and forms a critical salt bridge with D352, bisecting the nucleotide entry site in the closed conformation of the L2 loop. The presence of variant amino acid residues immediately upstream of K151 could have modulatory effects on the behaviour of the L2 loop. Clearly from our data, the D148A substitution has a substantial impact on viral RNA replication suggesting that it impairs the function of the L2 loop in the context of the HCVgt2a strain JFH1/Jc-1. By contrast, both double variants at positions 148 and 150 (D148Q.A150E) and a single variant at position 150 (A150V) are well tolerated. Thus, it is possible to introduce intragenotypic and intergenotypic substitutions into L2 without disrupting function. This indicates a degree of redundancy in the L2 loop sequences. However, it is possible that certain variants have subtle effects on the selectivity of incoming nucleotides. For example, the A150V variant gave reduced susceptibility to the nucleotide analogue sofosbuvir, which is a clinically approved and highly potent direct acting antiviral (15). This reduced potency could arise from lower capacity of the A150V substitution to allow sofosbuvir entry to the catalytic site of the polymerase. Furthermore, we see differences between the HCV subgenomic replicon and infectious virus assays for A150V (and P156A) such that these mutations affect viral replication in the replicon but not the HCVcc system. This may suggest differing roles of these sites in replication compared to assembly.

Our data reveal that mutations A150V and P156A influence viral replication in the presence of exogenous IFNλ4 such that, for both variants, the reduction in replication was less in both SGR and HCVcc systems compared to the WT control. These results suggest that V150 and A156 confer slight resistance to IFNλ4. The higher prevalence for V150 in those with *IFNL4* alleles that produce functional protein would suggest a fitness advantage for this variant, which may be consistent with the partial IFNλ4 resistance observed in our assays. However, viral load is lower in chronic infection in individuals who have *IFNL4* alleles that produce IFNλ4 compared to those who fail to make functional protein (2,3). Thus, our data illustrate the challenge with aligning *in vitro* results to understand the mechanisms underlying *in vivo* findings. It is possible that there are epistatic effects, which would be difficult to ascertain with *in vitro* methods that do not recapitulate infection by natural strains. In conclusion, we have characterised replicative effects of variants associated with *IFNL4* genotype that could serve as the basis for further studies into the role of viral variability in physiologically-relevant *in vitro* models.

## Author statements

CB and JM conceived the project and established the goals of the study; CB conducted all experiments; CB and JM analysed and interpreted the data; CB and JM were responsible for writing and preparing the manuscript.

## Acknowledgements

We wish to thank undergraduate students Maria Chalmers and Aqsa Sharif for their help with this project.

